# Vital signs analysis algorithm detects inflammatory response in premature infants with late onset sepsis and necrotizing enterocolitis

**DOI:** 10.1101/200329

**Authors:** Leena B. Mithal, Ram Yogev, Hannah L. Palac, Daniel Kaminsky, Ilan Gur, Karen K. Mestan

## Abstract

**Background:** Nonspecific clinical signs and suboptimal diagnostic tests limit accurate identification of late onset sepsis (LOS) and necrotizing enterocolitis (NEC) in premature infants, resulting in significant morbidity and antibiotic overuse. An infant’s systemic inflammatory response may be identified earlier than clinical suspicion through analysis of multiple vital signs by a computerized algorithm (RALIS).

**Aim:** To evaluate the revised RALIS algorithm for detection of LOS and NEC in preterm infants.

**Methods:** In this nested case-control study, VS data (heart rate, respiratory rate, temperature, desaturations, bradycardias) were extracted from medical records of infants 23-32 weeks gestation. RALIS generated an output, with score >5 triggering an alert. Patient episodes were classified based on culture, radiograph, and antibiotic data into categories: LOS, expanded LOS, NEC, and controls. Paired t-tests, linear regression and cross-validation analyses were used to evaluate the relationship between RALIS alert and LOS/NEC.

**Results:** Among 155 infants with 161 episodes, there were 41 expanded LOS (+ blood, CSF, urine, respiratory culture), 31 LOS (+ blood, CSF, urine), 9 NEC, and 93 controls. RALIS alert was 43.1+/-79 hours before culture in LOS (p=0.012). There was a significant association between RALIS alert and LOS/NEC (β=0.72, p<0.0001). Sensitivity and specificity for LOS/NEC were 84% and 80%, (PPV=63%; NPV=93%). The regression model demonstrated an AUC of 89.9%.

**Conclusions:** For infants <32 weeks, RALIS detects systemic inflammatory responses in LOS and NEC in the first month of life. The algorithm identifies infection earlier than clinical suspicion, even for NEC with negative cultures. RALIS has high NPV to rule-out LOS and NEC, and may, after prospective validation, aid in antibiotic treatment decisions.

## Introduction

Sepsis remains a critical issue and a major cause of death among infants in the U.S.(1) and causes approximately 200,000 annual neonatal deaths worldwide(2). Approximately 21% of very low birth weight (VLBW) infants are affected by at least one episode of late onset sepsis (LOS; with positive blood culture after 72 hours of life)(3). There has been some improvement in the incidence of LOS due to infection control measures(4), but LOS continues to disproportionately affect preterm infants and cause significant morbidities including neurologic impairment, prolonged hospitalization, and death(5). Because prompt recognition of infection and initiation of antibiotic therapy decrease sepsis-related morbidity and mortality, clinicians have a low threshold to evaluate for LOS and empirically treat with broad spectrum antibiotics. However, the lack of reliable diagnostic capability for LOS remains an ongoing issue in the neonatal intensive care unit (NICU). Clinical signs of sepsis are mostly nonspecific and detected late, and current laboratory tools are of limited utility. Specifically, the current gold standard for diagnosis of LOS is blood culture, which has delayed results and limited sensitivity in the setting of small specimen volumes and recent antibiotic administration(6). Adjunct markers of infection such as white blood cell count indices(7), C-reactive protein(8), hypoglycemia, and thrombocytopenia are also nonspecific with poor positive predictive value (PPV)(9, 10). Furthermore, there is lack of consensus on the definition of neonatal sepsis amongst providers(11). As a result, antibiotic treatment is overprescribed in the NICU with 56% of VLBW infants treated with antibiotics while only 21% had culture-proven infection in one large study(3). The adverse effects of prolonged antibiotic exposure in preterm infants include antibiotic resistance, fungal infections, necrotizing enterocolitis (NEC), and death(12–14). The nonspecific clinical presentation of NEC overlaps with that of sepsis. The early localized bowel inflammation, bacterial penetration, and tissue destruction is often difficult to identify until signs of abdominal distension, bloody stool, bowel perforation, or clinical deterioration appear, often accompanied by sepsis(15). Thus, development of more reliable and specific methods to predict and exclude LOS and/or NEC in preterm infants is essential to improve neonatal outcomes.

The advantages of automated vital signs monitoring as an objective tool to detect evolving sepsis is being increasingly recognized(16, 17). Several studies have suggested that analysis of vital signs patterns can help clinicians evaluate for impending infection or inflammatory response (e.g., sepsis, NEC) prior to obvious deterioration(18–21). For example, the HeRO monitoring system detects heart rate variability characteristics that change with LOS. The use of HeRO was associated with a reduction in 30 day mortality in VLBW infants(22, 23). The RALIS software was the first to incorporate multiple vital signs monitoring into a predictive algorithm to detect systemic inflammatory responses including LOS(24, 25). As we previously reported, RALIS detected sepsis 2.5 days prior to clinician suspicion of infection (defined as the time blood culture was obtained) with a sensitivity of 82% for LOS in infants ≤28 weeks gestational age (GA)(19). However, the PPV and negative predictive value (NPV) were only 67% and 65% respectively. Therefore, the RALIS algorithm was changed, primarily by incorporating more appropriate ranges for temperature and more accurate input of skin temperature rather than incubator values. The objective of the current study was to evaluate the performance of the revised RALIS algorithm to detect inflammatory responses from LOS and NEC in a cohort of preterm infants (23–32 weeks GA). Specifically, we compared time and frequency of RALIS alert to clinical suspicion of LOS among preterm infants with culture-confirmed LOS, NEC, and controls. Cross validation analysis was also performed to assess the robustness of the RALIS model.

## Materials and methods

### Patient population

Medical records of infants enrolled in a premature birth cohort study at Prentice Women’s Hospital and who were admitted to the NICU were reviewed. Per standard study protocol, parental consent is obtained, patients are assigned a study code, and a de-identified database is stored on a secure server. Included infants were born at <33 weeks GA between 2008 and 2012 with complete VS data from birth to 28 days of life available in the electronic medical record. Infants transferred out of the NICU, who died within the first month of life, and with congenital syndromes were excluded. Infants requiring oscillator or jet ventilation were also excluded because of inability to interpret respiratory rates. Maternal and infant characteristics including demographics and clinical course were collected using the cohort database, Northwestern University Enterprise Data Warehouse system, and standardized medical record review. Extracted data included comprehensive microbiologic culture information, type and duration of antibiotics, and laboratory results. This study was approved by the Institutional Review Board of Northwestern University and Ann and Robert H. Lurie Children’s Hospital of Chicago.

### Vital signs and RALIS output

Vital signs (VS) data are entered into the electronic medical record (PowerChart, Cerner, MO) by experienced nurses per routine NICU protocol. Nurses read and record the VS from the standard continuous cardiorespiratory monitors (IntelliVue Neonatal, Philips, MA). All patients had vital signs documented at least every 3 hours for 28 days of hospital admission. The VS data was extracted retrospectively, coded in Excel, and uploaded into the RALIS program in order to generate continuous RALIS output. RALIS is a mathematical algorithm developed for monitoring of preterm infants to detect inflammatory response, such as in LOS. As previously described, VS data incorporated into RALIS include: heart rate, respiratory rate, temperature, desaturation events, and bradycardia events(19). The heart rate, respiratory rate, and temperature values are numerical, while the low SpO2 (oxygen saturation) and bradycardic inputs are binary, indicating the presence or absence of events during the preceding 2-3 hour interval. A desaturation event was defined as an oxygen saturation of <80% for >10 seconds. A bradycardia event was defined as a heart rate <100 beats per minute for >10 seconds.

The software generates a RALIS score based on significant VS changes from the individual patient’s baseline, which is initially calibrated from the first 72 hours of life and evolves over the monitoring period. Additionally there are GA and birth weight specific VS ranges incorporated into the algorithm. Weight is entered once every 24 hours. The final RALIS score is reported on a 0-10 scale, with 5 as the threshold for an acute inflammatory response based on an initial derivation cohort in Israel. A RALIS score ≥5 for 6 consecutive hours generates an alert to the clinical team. We limited our analysis to LOS and NEC episodes within 3-28 days of life for consistency and comparison with previous studies(19, 25). A RALIS alert was considered associated with a LOS or NEC episode if it occurred within 7 days before or after infection was suspected.

### LOS definitions

The primary outcome: LOS was defined as an episode (physician-initiated sepsis evaluation) occurring at >72 hours of life which resulted in a positive culture and was treated with an antibiotic treatment course. A positive culture was defined as any blood, cerebrospinal fluid (CSF), or urine culture with bacteria or yeast. We also collected data on episodes with a positive endotracheal tube culture (ETT) and chest radiograph infiltrate treated with antibiotics: which were included in the *expanded* LOS classification. Antibiotic treatment was receipt of a ≥7 day of antibiotic course. Neonatal complications such as NEC(26) and bronchopulmonary dysplasia (BPD)(27) were identified according to parent study protocol and widely used benchmarks. Specifically, an episode of NEC was defined as clinical and radiographic evidence of NEC (pneumatosis intestinalis) treated with antibiotics and bowel rest. Based on NICHD definitions for premature infants, episodes of culture-proven sepsis occurring <72 hours of life were considered early onset and excluded from analysis. Patients with no positive culture and no antibiotic course throughout their hospitalization served as controls. Clinical sepsis patient episodes with negative culture, but that received antibiotic treatment were classified as culture-negative LOS. A positive culture without antibiotic treatment in an infant who did clinically well was considered a false-positive culture episode.

### Statistics and Modeling

For sample size determination, we considered LOS the primary outcome and the hours between RALIS alert and culture as the primary predictor. Based on our previous study, we used a mean time difference of 59 (±67) hours in the LOS group and 22 (±64) in the false-positive culture group (as a proxy for no LOS)(19). Thus, our cohort was adequate with greater than the estimated sample size of 24 LOS cases to achieve >80% power (alpha 0.05). Statistical analysis was performed by using SAS^®^ 9.4 (Cary, NC) and Stata^®^/IC 13.1 (College Station, TX).

Demographic and clinical characteristics are reported as means with standard deviations and frequencies with percentages. Differences in characteristics among patients were assessed using independent samples t-tests, chi-square, or Fisher’s exact tests. The association of RALIS with sepsis episodes was evaluated by comparing the hours between the time of RALIS alert and the time of clinical suspicion of sepsis, defined as the time a specimen was sent for culture. Time differences between RALIS and clinical suspicion were compared using the Wilcoxon signed-rank test. Linear regression was used to evaluate the relationship between the timing of RALIS alert and culture. We calculated sensitivity, specificity, PPV and NPV of RALIS for 1) expanded LOS (including blood, CSF, urine, and ETT positive cultures), 2) LOS (including only blood, CSF, and urine positive cultures), 3) NEC, and 4) LOS (blood, CSF, urine) plus NEC. All episodes were treated independently. Two-sided p-values <0.05 were considered statistically significant.

We implemented an *exploratory* cross-validation analysis using a modified hold out method to assess the validity of RALIS alert to predict LOS using logistic regression modeling. Patients with true positive (expanded LOS) and true negative status for infection (controls) were randomly assigned to the train dataset (75%) or the test dataset (25%). The train dataset was used to fit a multiple logistic regression model using a manual stepwise selection procedure with a model entry criterion of p=0.10. Patients in the test dataset were not included in model development. To assess the predictive value of this model, we evaluated the area under the curve (AUC) of the receiver operating characteristic (ROC) curve in the train and test datasets. The test dataset was used to cross-validate the model by comparing the R^2^ of the predicted probabilities of infection and model-estimated coefficients for RALIS alert to those from the train dataset. A relative percent difference of 10% between the train and test model-estimated coefficients were chosen *a priori* as the validation threshold.

## Results

### Demographics

The 155 patients had 161 episodes including: 41 cases of expanded LOS, 14 culture-negative sepsis, 13 false-positive cultures, and 93 controls. Of the LOS positive cultures: 22 were blood, 8 urine, 10 ETT, and 1 blood+CSF. Organisms were gram negative rods (n=10), *S. aureus* (n=8), coagulase negative Staphylococcus species (CONS, n=13), group B Streptococcus (n=1), Enterococcus species (n=6), Candida (n=1), and 2 polymicrobial episodes with a gram negative rod and *E. faecalis* or *S. aureus.* There were 9 cases of NEC within the first 28 days of life (2 concurrent with blood culture-proven LOS episodes and 7 with culture-negative LOS episodes). Thus, the total episodes of LOS and/or NEC was 38.

As can be seen in Table 1, the control group had a significantly higher gestational age (30 (1.7)) weeks compared to the expanded LOS group (26.7 (2.3) weeks), the LOS group (27.2 (2.3) weeks), and the false-positive culture group (27.4 (1.1) weeks). The expanded LOS and LOS categorizations (with and without inclusion of ETT cultures n=41 and 31 respectively) had no notable differences in demographics. Controls also had a higher birthweight and lower incidence of BPD than other groups. Compared to controls, chorioamnionitis was more prevalent in the expanded LOS and LOS groups. Gender, multiple gestation, and prolonged rupture of membranes were not significantly different amongst groups.

**Table 1.**
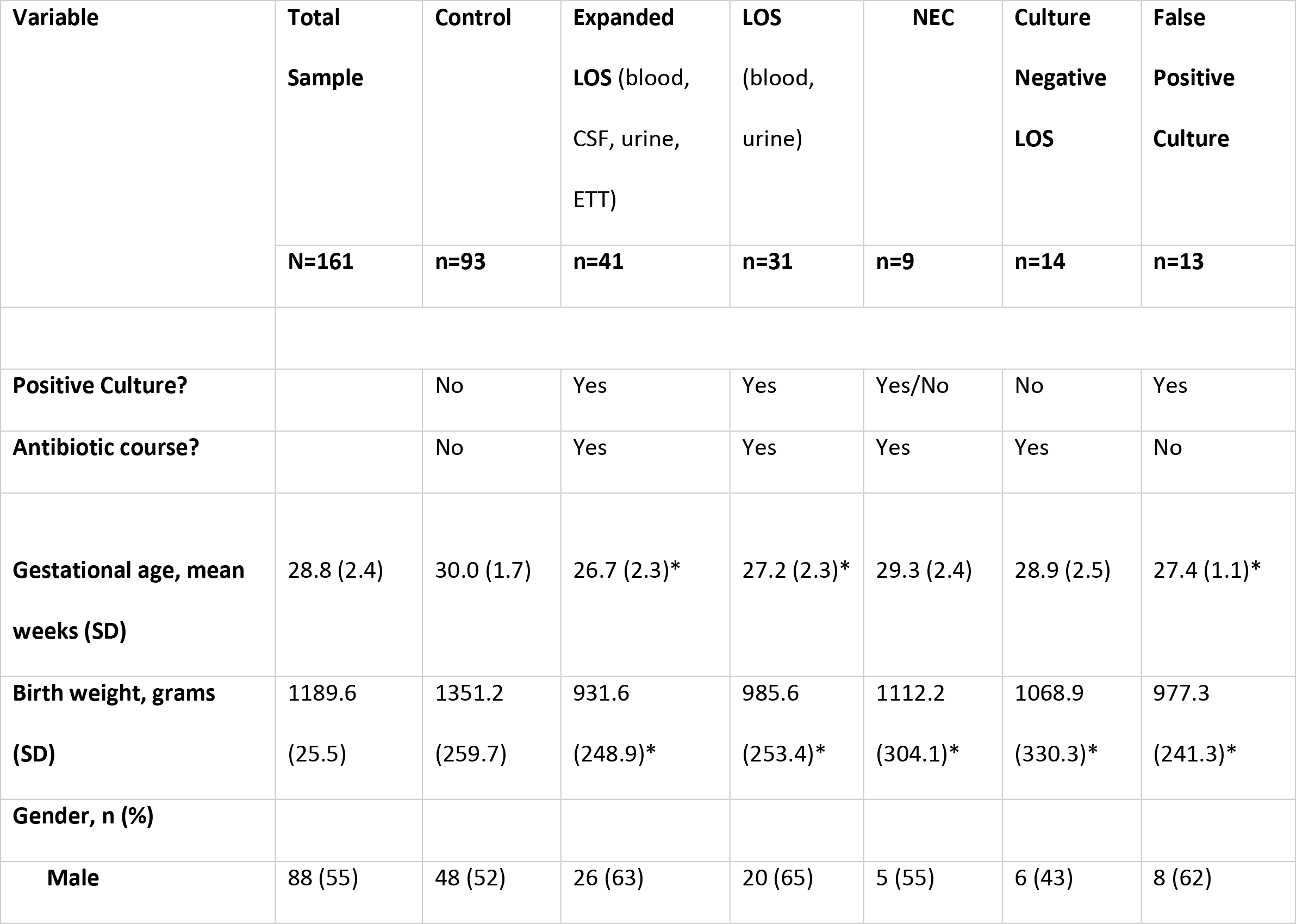
Demographic and Clinical Characteristics of the Patient Sample±.

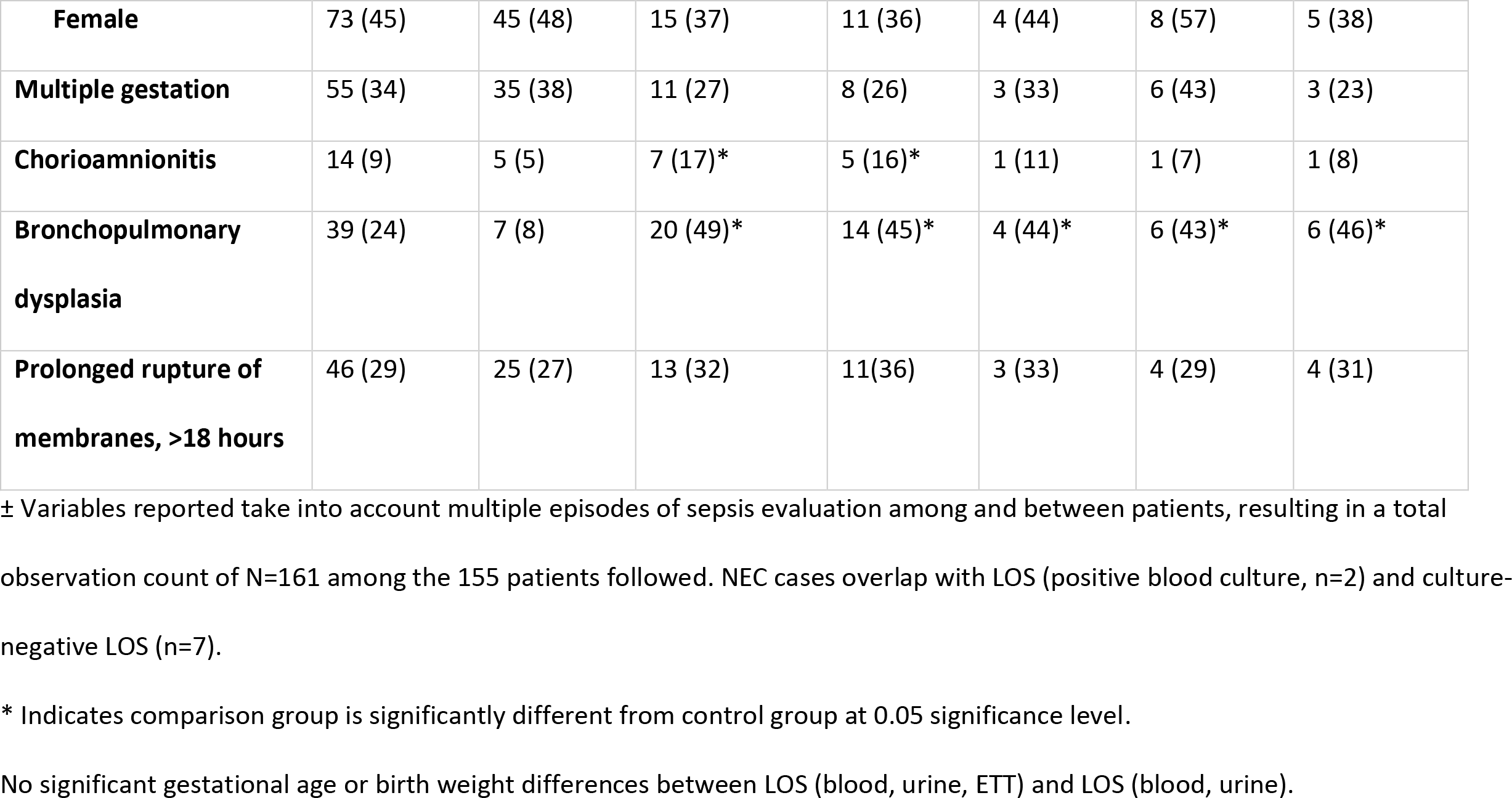

### RALIS alert

RALIS had an associated alert in 25 of 31 LOS episodes (81%). Of the 6 LOS episodes without an associated alert, 4 were CONS bacteremia. In the control patients, 74 of 93 had no RALIS alert (80%), and 19 (20%) had a false-positive alert (Table 2). The sensitivity of RALIS alert for LOS was 81% and specificity was 80%, with a PPV of 57% and NPV of 93%. For expanded LOS, the sensitivity was 84%, with a PPV of 63% and NPV of 93%. Amongst the 9 cases of NEC, 7 of 7 culture-negative cases had an associated RALIS alert (100%). Two episodes of NEC occurred with a concurrent blood-culture proven LOS episode, of which 1 had an associated alert. The sensitivity of RALIS alert for LOS and/or NEC (n=38) compared to controls (n=93) was 84%, with specificity of 80%, PPV of 63%, and NPV of 93%. Eleven of 14 culture-negative sepsis episodes had an associated RALIS alert (79%).

**Table 2.**
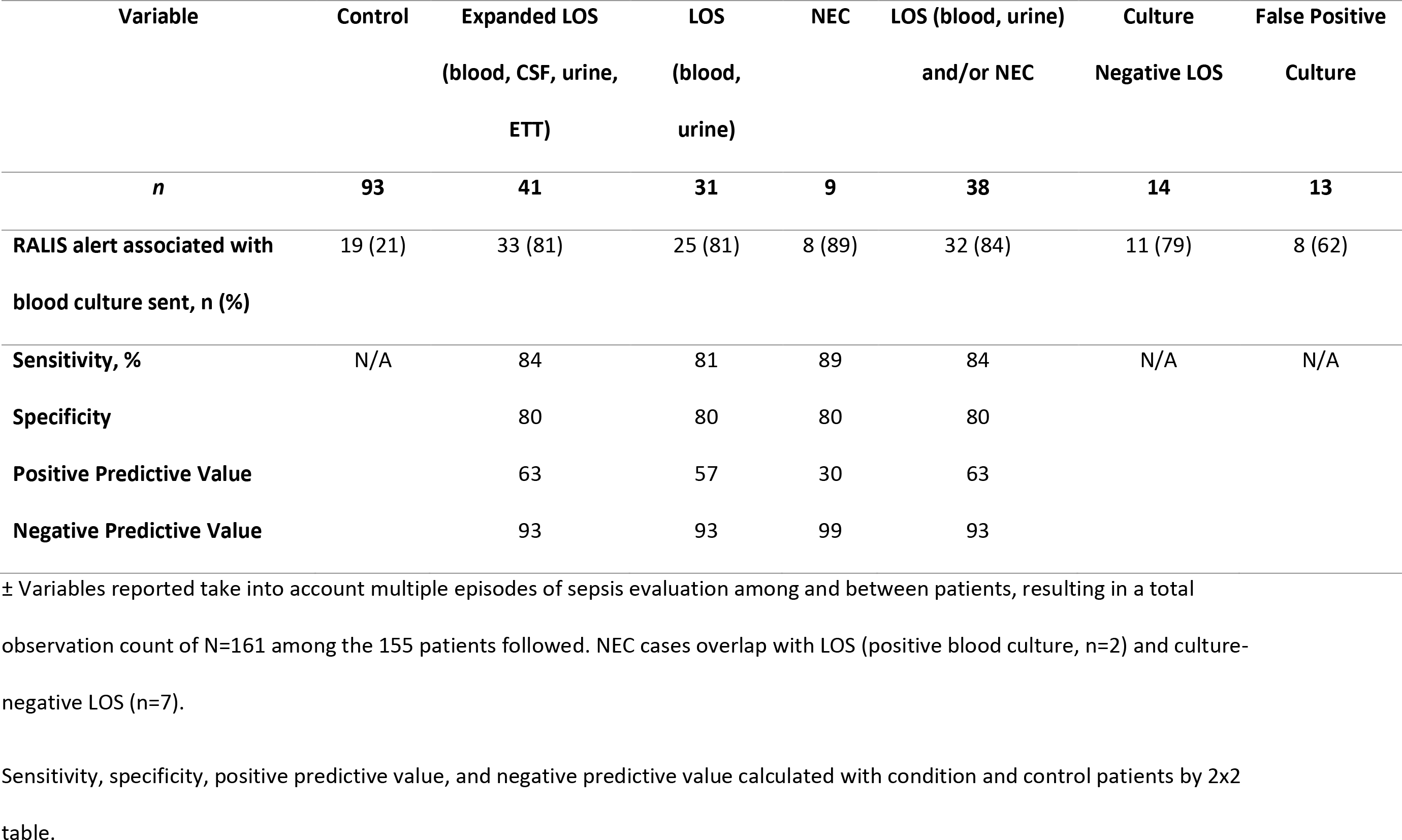
RALIS output±.

For the LOS group, the mean hours between RALIS alert and culture sent was −43.1 ±79.4 hours [mean±SD] (p=0.012, 95% CI: −75.8 to −10.3 hours). Linear regression analysis showed a significant association between RALIS alert time and culture time (β-coefficient =0.72, p<0.0001). The same statistical analysis was conducted for LOS and/or NEC as the outcome. Again, mean time of alert was earlier, −33.0±79.3 hours before culture (p=0.025, 95% CI: −61.5 to −4.4 hours). There was a significant linear association between RALIS alert time and LOS/NEC (β-coefficient =0.72, p<0.0001) (Figure 1).

**Figure 1.**
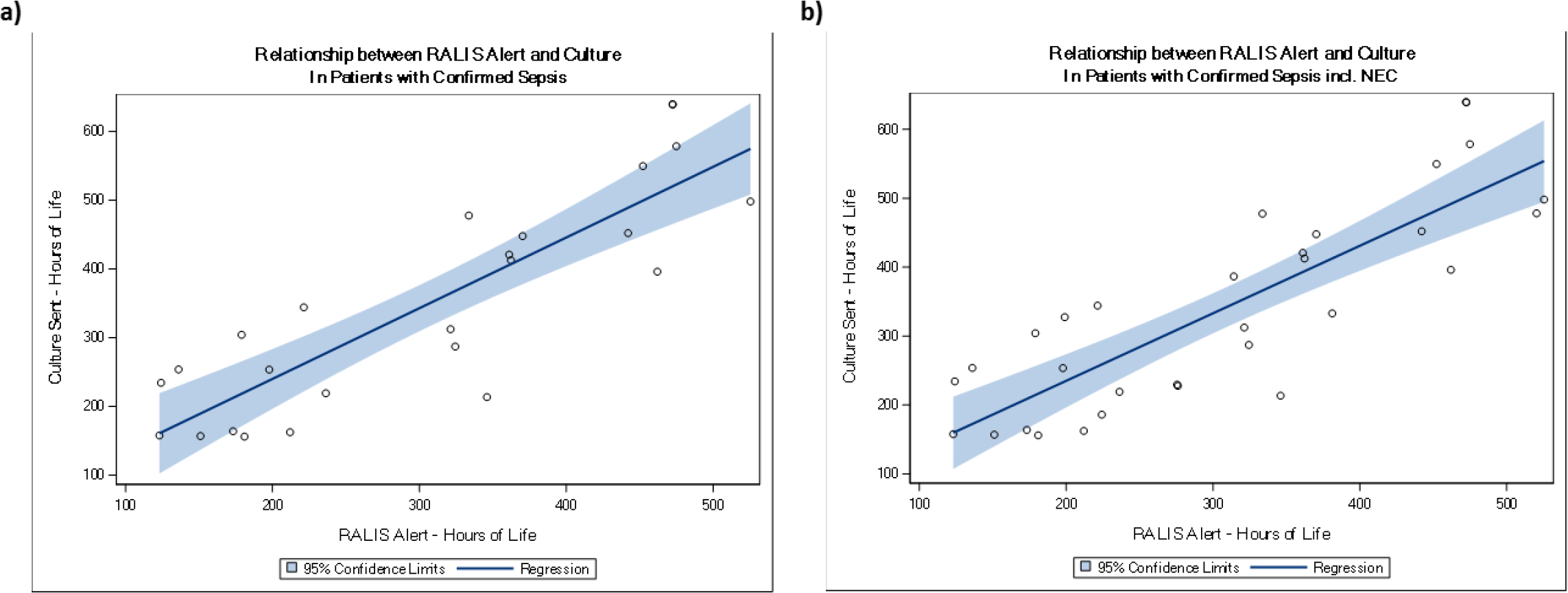
Linear Regression Plots of RALIS alert with Culture. (a) The regression line for the bivariate association between culture hours of life (HOL) and RALIS alert HOL for LOS [blood, urine]: RALIS Alert HOL − 54.2+ 0.72*Culture HOL. b) Regression line equation for LOS and/or NEC: RALIS Alert HOL − 62.9 + 0.72 *Culture HOL. The shaded region represents the 95% confidence interval for the regression line.

### Cross validation

The expanded LOS and control patients were randomly divided into the train dataset (n=100, LOS cases=29) and test dataset (n=34, LOS cases=12). Birthweight (p=0.0035) was included in the model, while GA, gender, multiple gestation, chorioamnionitis, and prolonged rupture of membranes were not significant as predictors of infection. The AUC of the final logistic regression model in the training dataset was 89.9% [95% CI: 82.5-97.2%] (Figure 2). The model-estimated coefficient for RALIS alert in the training dataset was 1.7250; the model-estimated coefficient in the test dataset was 1.8276. The R^2^ of the model was 0.3811 in the training dataset and 0.4598 in the test dataset, with an absolute change (increase) in R^2^ of 0.0790 and a 5.95% relative percent change in model-estimated coefficients.

**Figure 2.**
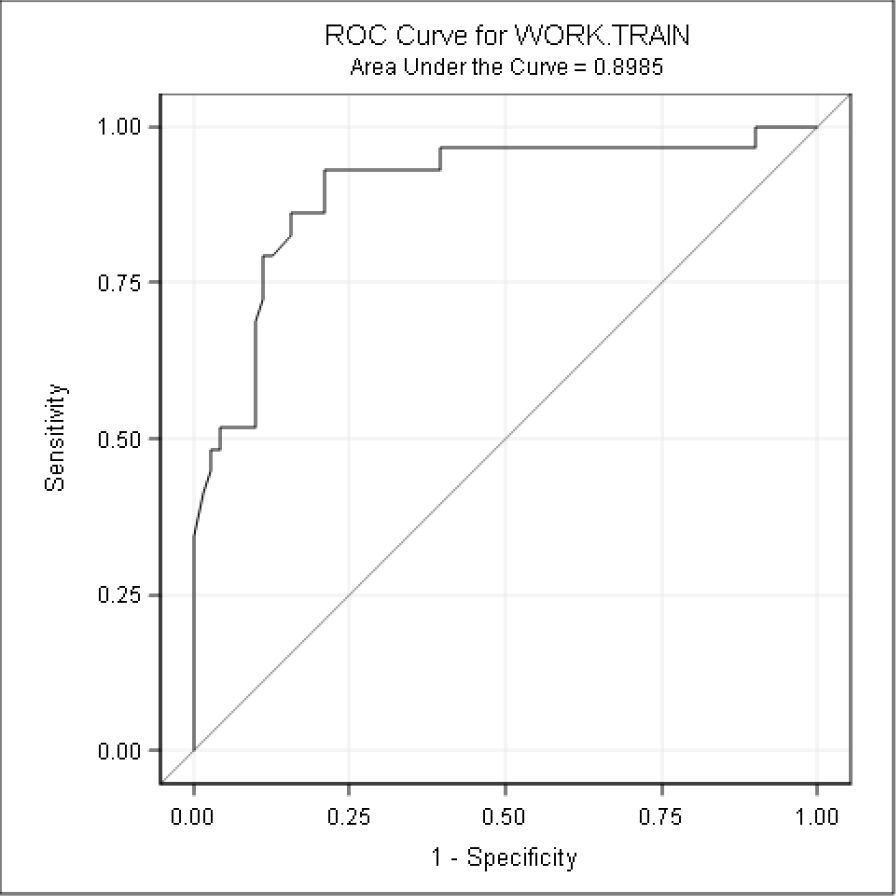
ROC Curve for Logistic Regression Model in Cross Validation Analysis. The AUC of the final derivation model in the testing dataset using model parameter estimates yielded an AUC of 0.8985.

## Discussion

We have previously tested the RALIS multiple vital signs analysis algorithm in preterm infants for detection of inflammatory response in the setting of LOS. We expanded our work in this study to test an improved version of the RALIS software in a broader GA range and included NEC. Our major finding is that the revised RALIS algorithm functions well in detection of LOS and NEC inflammatory responses within the first four weeks of life, and has particular utility in the exclusion of LOS and NEC with a high NPV. There was an associated alert in the vast majority of NEC episodes, even with negative blood cultures. On average, a RALIS alert occurred significantly earlier than clinical suspicion of LOS or NEC. Though limited in sample size, in exploratory cross validation analysis, the model estimated coefficients for RALIS in the train and test models remained similar. This reinforces the model of RALIS alert with birthweight in predicting inflammatory responses of LOS and NEC in premature infants ≤32 weeks.

The reliable detection and exclusion of LOS and NEC are ongoing clinical challenges in the NICU, given limited diagnostic tools. Thus, vital signs analysis can be a useful, additive approach. The HeRO system has demonstrated signals in LOS approximately 24 hours prior to clinical diagnosis(21). In a subsequent study, Fairchild and colleagues reported a cross-correlation between heart rate and oxygen saturation, demonstrating the potential for other vital signs to improve HRC index monitoring for detection of LOS(28). There is evidence that acute increases in apnea, bradycardia, and desaturation episodes are some of the most common signs of LOS in preterm infants in the NICU(29). Thus, incorporation of multiple vital signs, including episodes of bradycardia and desaturations, as indicators of the pathophysiologic response to acute systemic inflammatory response is logical and innovative.

RALIS is based on this multifaceted approach and has now been tested through iterations of revised algorithms to maximize its predictive value (both positive and negative) for clinical use in the NICU. RALIS overcomes the issue of significant interindividual variation in VS by using GA and birthweight specific ranges in combination with acute changes from an individual infant’s own baseline. Our current data demonstrates an 84% sensitivity of RALIS for inflammatory response in LOS and/or NEC with a high NPV of 93%. It is notable that of the cases of false-negative RALIS output (cases of LOS with no associated alert n=6), two-thirds were CONS species in the blood (n=4), an organism that typically does not elicit a robust systemic inflammatory response. It is also possible that these 4 episodes were reflective of culture contaminants and may not have required antibiotic treatment.

RALIS can detect LOS and NEC before a potential decompensation with mean alerts 33 hours prior to clinician suspicion, but perhaps more importantly, it may reassure a clinician of the absence of LOS and NEC by displaying a normal score. By providing objective evidence that systemic inflammatory signs are not present, clinicians may more confidently withhold or discontinue antibiotics, thus potentially reducing the burden of empiric antibiotic overuse in preterm infants. Early life antibiotic exposures not only cause short term complications during the NICU hospitalization, but there is growing literature about the far-reaching effects of antibiotics on risk of chronic conditions such as obesity and asthma later in life(30–32). Antibiotic use for LOS in the NICU remains highly variable and clinicians could benefit from additional evidence-based data(33, 34). The PPV of a RALIS alert is similar to that reported in our previous study (67% vs 63%)(19), with a notably higher NPV (65% vs 93%). The only change in the study subjects was an expansion of gestational age. All other exclusion criteria (oscillator ventilation, proven EOS, transfer or death within first month of life) remained the same between our previous and current study. The exclusion of these infants, who were potentially more severely ill, is a limitation to general applicability of this technology at this point. RALIS monitoring for critically ill infants requiring oscillator ventilation and with EOS requires further study. Alternative approaches may need to be developed to incorporate respiratory rate in these patients at high risk for negative outcomes. Of note, there were also some false-positive alerts in control patients that would trigger clinicians to evaluate the patient and likely send diagnostic tests for infection. The balance between additional diagnostic laboratory tests to rule-out sepsis versus discontinuation of antibiotics attributable to use of the RALIS algorithm requires further prospective studies.

Sample size constraints remain a limitation in this study, particularly for cross-validation analysis, as is the retrospective nature of the study in a single center. Infants in the control group had significantly higher GA and birth weight than LOS and NEC groups. This is a source of potential bias, and thus future studies may require a frequency matching approach with these parameters. The strengths of this study include 1) detailed review of the patient’s clinical data and precise definitions of culture-proven LOS (with clear delineation of culture source) and 2) removal of both culture-negative LOS and false-positive culture cases from regression analysis, because clinical definitions of LOS are biased by variable physician antibiotic prescribing practices. The inclusion of NEC adds to the confidence in RALIS for detection of systemic inflammatory response from an early evolving infection. In the future, the utility of RALIS for detection of inflammatory response in LOS and NEC prior to clinical suspicion and for its reliable negative signal requires confirmation and further study in a larger, multi-center prospective trial. In addition, RALIS monitoring needs to be investigated for its utility after the first month of life.

## Conclusions

The revised RALIS algorithm of multiple VS analysis can detect acute subclinical changes indicative of systemic inflammatory response such as in LOS and NEC. This retrospective study provides evidence that the RALIS alert identifies preterm infants with developing infection in the first month of life earlier than suspected by a clinician and appears to do so even in cases of NEC with negative cultures. The high NPV suggests RALIS may be particularly useful in ruling out LOS and NEC, allowing discontinuation of unnecessary antibiotics in preterm infants who are at highest risk of adverse effects from antibiotic overuse. RALIS may aid in medical decision-making in combination with other clinical and laboratory assessment. Thus, the real time application of this tool is promising and warrants further evaluation in prospective trials.

## Acknowledgements

The authors would like to thank Moheet Merchant for his assistance in vital signs data extraction. This work was supported by the National Institutes of Health [National Heart, Lung, and Blood Institute K23 HL093302] and the Northwestern Memorial Foundation Friends of Prentice Grants Initiative.

